# Substantial hierarchical reductions of genetic and morphological traits in the evolution of rotiferan parasites

**DOI:** 10.1101/2024.08.01.605096

**Authors:** Holger Herlyn, Anju Angelina Hembrom, Juan-Pablo Tosar, Katharina M. Mauer, Hanno Schmidt, Bahram Sayyaf Dezfuli, Thomas Hankeln, Lutz Bachmann, Peter Sarkies, Kevin J. Peterson, Bastian Fromm

## Abstract

Graphical abstract

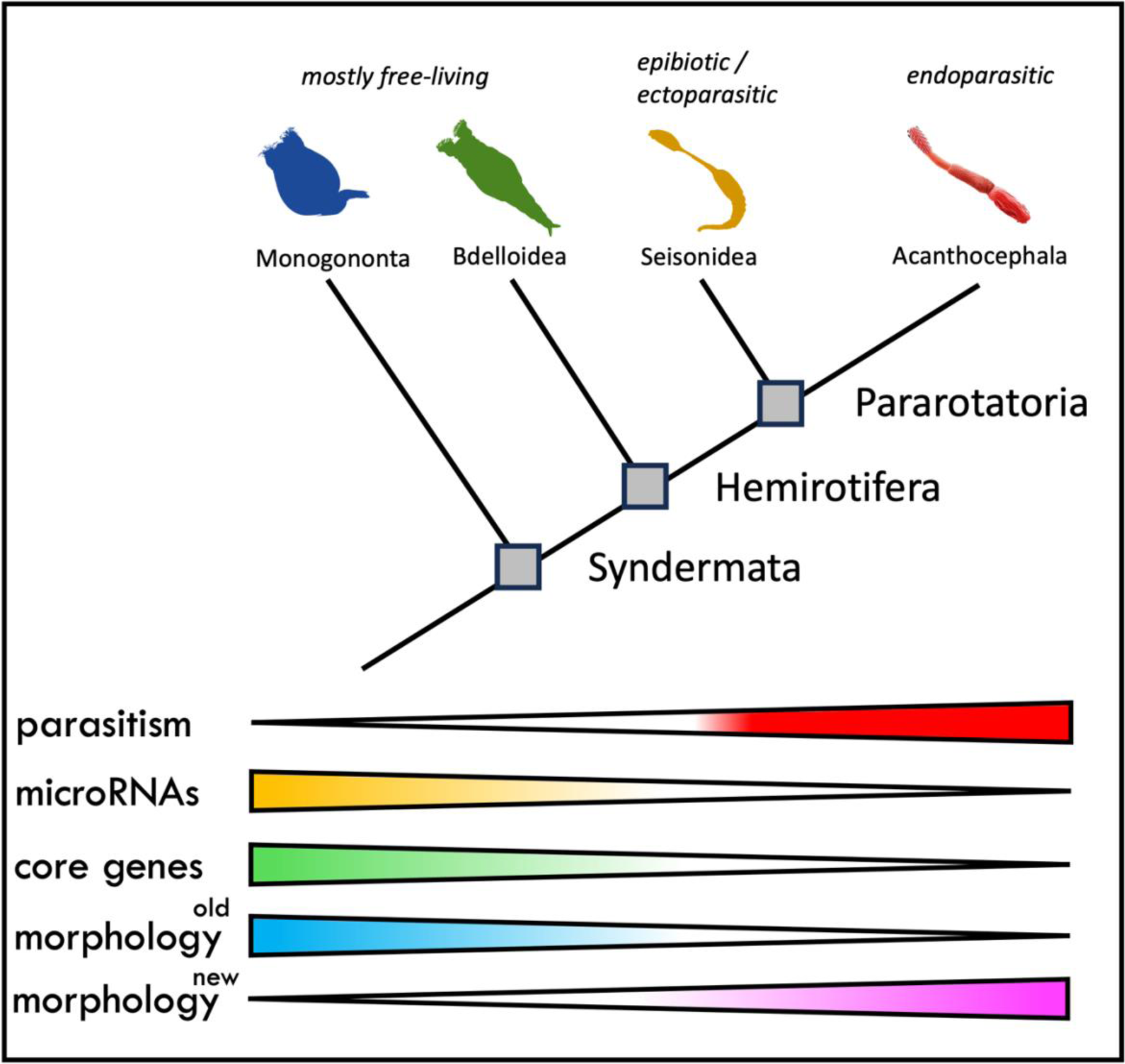

During the last 800 million years of evolution animals radiated into a vast range of diversity of species and disparity of forms and lifestyles. The process involved a near hierarchical increase in complexity from life forms with few cell types to organisms with many hundreds of cell-types. However, neither genome size nor number of protein-coding genes can explain these differences and their biological basis remains elusive. Yet, recent studies have suggested that the evolution of complexity is closely linked to the acquisition of a class of protein coding gene-regulators called microRNAs.

In a regressive approach, to investigate the association between loss of organismal complexity and microRNAs, we here studied Syndermata, an invertebrate group including free-living rotifers (Monogononta, Bdelloidea), the epibiotic Seisonidea and the endoparasitic Acanthocephala. Genomic, transcriptomic and morphological characterization and comparisons across 25 syndermatan species revealed a strong correlation between loss of microRNAs, loss of protein-coding genes and decreasing morphological complexity. The near hierarchical loss extends to ∼85% loss of microRNAs and a ∼50% loss of BUSCO genes in the endoparasitic Acanthocephala, the most reduced group we studied.

Together, the loss of ∼400 protein-coding genes and ∼10 metazoan core gene losses went along with one microRNA family loss. Furthermore, the loss of ∼4 microRNA families or ∼34 metazoan core genes associated with one lost morphological feature. These are the first quantitative insights into the regulatory impact of microRNAs on organismic complexity as a predictable consequence in regressive evolution of parasites.

## Introduction

The evolution from life forms with very few cell types, to more complex organisms with many cell-types, tissues and organs, and substantial cognitive abilities must have a genetic basis (Valentine et al. 1994). However, the detailed nature of the complexity-genome relation remains to be clarified, since neither differences in genome size (Cavalier-Smith 1978; Gregory 2001), nor in the number of protein-coding genes (Hahn and Wray 2002) are indicative of animal complexity. In fact, the molecular basis of the large divergence in organismic complexity remains one of the biggest mysteries in biology.

Several recent studies suggested that the evolution of animal complexity, novel cell types and tissue identities is hierarchical and tightly correlated with the acquisition of microRNAs (Sempere et al. 2006; Tanzer and Stadler 2006; Grimson et al. 2008; Heimberg et al. 2008; Peterson et al. 2009; Wheeler et al. 2009; Christodoulou et al. 2010; Berezikov 2011; Fromm et al. 2015; Alberti et al. 2018; Deline et al. 2018; Zolotarov et al. 2022; Fromm 2024). microRNAs are ∼22 nucleotide short non-coding gene-regulators involved in a plethora of biological processes (Bartel 2018). By regulating messenger RNAs translation, either in a switch-like or rheostat manner, microRNAs can diversify existing genetic programs and can canalize the evolution of new cell types and phenotypes (Hornstein and Shomron 2006; Peterson et al. 2009; Wu et al. 2009; Avital et al. 2018; Chakraborty et al. 2020). Their importance is reflected in deep evolutionary conservation and rare losses from genomes making them excellent phylogenetic and taxonomic markers ((Wheeler et al. 2009; Helm et al. 2012; Tarver et al. 2013; Kenny et al. 2015; Tarver et al. 2018; Fromm et al. 2019; Fromm 2024) down to the species level (Fromm et al. 2014; Ovchinnikov et al. 2017; Kang et al. 2018). However, due to large evolutionary timescales, direct observation of increases in complexity due to microRNA expansion are difficult (but see (Jenike et al. 2023)). If microRNAs are indeed key players in animal complexity, then the secondary reduction of morphological features in animals transitioning from free-living to parasitic lifestyles presents a unique opportunity to test the microRNA-to-complexity correlation: microRNA repertoires should decrease in conjunction with simplified morphology.

Parasitism has evolved more than 200 times independently in animals since as early as the Cambrian (Weinstein and Kuris 2016; Cong et al. 2017). The co-evolutionary arms race between parasites and their hosts substantially contributed to the large diversity of species living today. There are an estimated ∼300,000 parasitic helminth species alone, infecting nearly all of the ∼79,000 known vertebrate species (Dobson et al. 2008; Parr et al. 2014). The adaptations to live on or inside hosts is usually accompanied by substantial changes to morphology, so that many parasites hardly resemble their free-living relatives and show striking reductions in organismic complexity, e.g., through losses of whole organs or the development of new structures for attachment. More recently, it has become clear that parasite evolution is often reflected in genome simplification, including large losses of genes and regulatory elements. However, it is highly debated whether morphological reductions in parasite evolution always go along with loss of genomic features or whether the two can occur independently (Tsai et al. 2013; Hahn et al. 2014; Zarowiecki and Berriman 2015; International Helminth Genomes Consortium 2019) (and see (Jackson 2015; Adams et al. 2020)).

We addressed whether there is a connection between the reduction of complexity and microRNAs in Syndermata AHLRICHS, 1997. The protostome group is composed of the microscopic ‘wheel animals’ (Rotifera CUVIER, 1817) belonging to either Monogononta PLATE, 1889 or Bdelloidea HUDSON, 1884, as well as Pararotatoria (Sielaff et al. 2016; Vasilikopoulos et al. 2024). The latter includes Seisonidea WESENBERG-LUND, 1899 measuring in the millimeter range, and macroscopic ‘thorny-headed worms’ or Acanthocephala KOELREUTER, 1771. Most monogononts and bdelloids are free-living, while the transition to a parasitic life-style occurred rarely in both taxa (May 1989). Moreover, seisonids are epibionts living strictly on leptostracan crustaceans (possibly as ectoparasites (Plate 1887)). Lastly, endoparasitic acanthocephalans exploit mandibulates as intermediate and gnathostome vertebrates as definitive hosts (Meyer 1933; Herlyn 2021).

We analyzed syndermatans for their microRNA complements, protein-coding gene repertoires and morphology. For this purpose, we generated the first smallRNA sequencing datasets for two acanthocephalan and one seisonid species. Together with publicly available genome and smallRNA sequencing data, we annotated microRNA complements of 25 syndermatan species including 11 monogonont and 11 bdelloid rotifers as well as the one seisonid and two acanthocephalan representatives. We further compared sizes of protein-coding gene complements, analyzed single copy protein-coding core gene complements (BUSCO) (Manni et al. 2021) and incorporated an extended morphological character matrix. Within Syndermata, we found a near-hierarchical loss of microRNAs from free-living rotifers to epibiotic seisonids and endoparasitic acanthocephalans, with 25% of the expected microRNA families missing in monogononts and bdelloids, and an unprecedented overall loss of 67% and 85% of usually conserved microRNA families in the seisonid and acanthocephalan lineage, respectively. We found the substantial loss of microRNAs to be highly correlated with a likewise strong loss of protein coding genes numbers and similarly hierarchical reductions in the highly conserved core complements (BUSCO). Losses of microRNAs and protein coding genes correlated strongly with the loss of morphological features.

Together, this study reveals that the secondary loss of complexity in Syndermata from free-living to epibiotic and endoparasitic groups is hierarchical and associated with substantial reductions in the complements of microRNAs and protein-coding genes, supporting the hypothesis that microRNAs are closely intertwined with organismal complexity.

## Results

### The microRNA complements of syndermatans show drastic hierarchical losses

Given their position within Metazoa (e.g. (Struck et al. 2014; Bleidorn 2019; Laumer et al. 2019)), 1 eumetazoan, 31 bilaterian and 12 protostome microRNA families were expected to be conserved in syndermatans. However, significant numbers of the corresponding 44 microRNA families were lacking in the species studied (Figure 1 A & B) according to genome-based predictions by MirMachine (Umu et al. 2023). The losses amounted to at least 10 microRNA families in free-living wheel-animals (Bdelloidea: 10 losses, Monogononta: 14 losses). The epibiotic representative of Seisonidea, *Seison nebaliae*, lacked 30 of the expected microRNA families, and 37 microRNA families were absent in endoparasitic Acanthocephala (Figure 1A). Complementary analyses of smallRNA sequencing datasets with MirMiner (Wheeler et al. 2009) identified six additional microRNA families that potentially emerged in the syndermatan stem line. Corresponding genes were determined in the monogonont and bdelloid representatives whereas only three and one occurred in the seisonid and the acanthocephalan species, respectively (Figure 1A). For the other microRNA families, MirMiner confirmed paralogue numbers as predicted by MirMachine. This included the identification of additional gene copies for most microRNA families in *Adineta vaga*, reflecting genome duplication in the bdelloid stem line (Simion et al. 2021). Together, MirMachine and MirMiner annotations yielded 54 microRNAs for the monogonont *Brachionus plicatilis* (35 conserved families), 129 microRNAs for the bdelloid *A. vaga* (39 conserved families), and only 29 microRNAs for the seisonid *S. nebaliae* (16 conserved families). Even fewer microRNAs were identified in the acanthocephalans, with 15 (7 conserved families) in *Neoechinorhynchus agilis* and 12 (7 conserved families) in *Pomphorhynchus laevis*. Relative to the expectation, microRNA family losses amounted to 20% in bdelloids, 29% in monogononts, 67% in the seisonid and 85% in the acanthocephalans. This is suggestive of hierarchical and progressive reduction of microRNA repertoires in the lineage to Acanthocephala (Figure 1B), whereby the levels reached in thorny-headed worms are the lowest reported for any bilaterian animal to date. MirMiner additionally predicted several novel species-specific microRNAs (Figure 1B). Novel and conserved microRNAs displayed expected hairpin lengths of ∼60 bp in monogononts, bdelloids and acanthocephalans while the seisonid, *S. nebaliae*, was distinguished by remarkably long hairpins of more than ∼200 bp. Despite dramatic differences in the length of pre-microRNAs, mature sequences showed the expected high level of conservation (Tarver et al. 2013) as exemplified in MIR-1 (Figure 1D,E).

**Figure 1:**
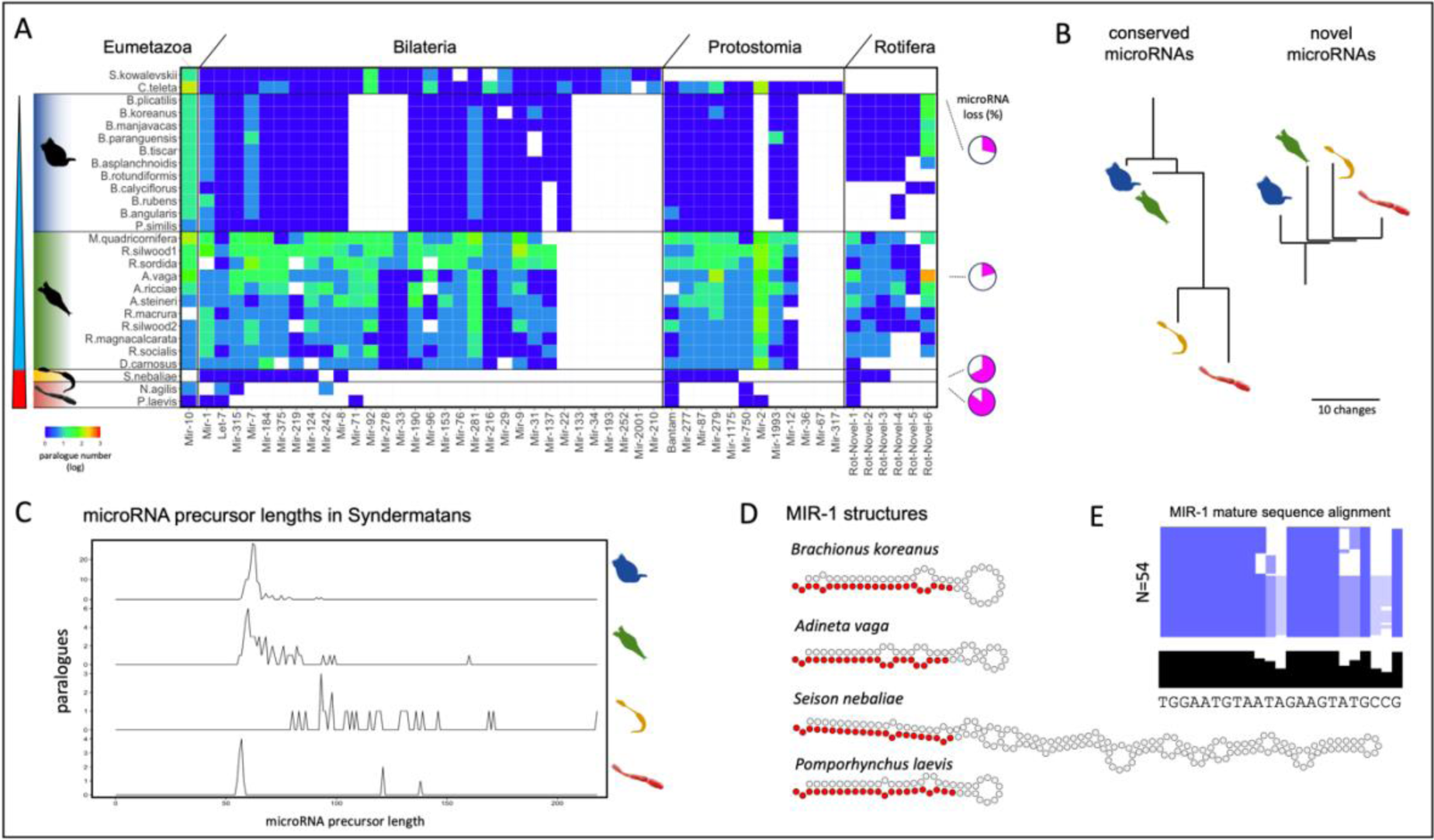
The microRNA complement of Syndermata shows stepwise losses from free-living to parasitic groups and unprecedented losses of 85% of conserved microRNAs in acanthocephalans. A) Banner-plot of 25 microRNA complements of included syndermatan species and two outgroups. White field means that no microRNA was found. Heatmap function refers to the paralogue number in each microRNA family. Note that both outgroups are not Syndermatans, and, hence, have none of the Rotifera microRNA families, and *S. kowalevskii* is not a protostome, and hence has no protostome microRNA families. B) Schematic trees of the four syndermatan groups representing loss of conserved and gain of novel microRNA families. Branch lengths correspond to the number of gains and losses. C) Overview of microRNA precursor lengths in selected representatives of monogononts (*B. plicatilis*), bdelloids (*A. vaga*), the seisonid *S. nebaliae* and the acanthocephalan *P. laevis*. D) Selected MIR-1 examples of the same species as in C) highlighting the length deviations in *S. nebaliae*. E) Alignment of the mature sequence of all MIR-1 genes in syndermatans (N=54) highlights its very strong conservation in the pattern characterized for bilaterian mature miRNAs (Wheeler et al. 2009; Fromm et al. 2015) with strong conservation in the seed and 3’ CR and higher variability in positions 9-12 and 17-22. Species order corresponds to that in Figure 1A, consensus in black.

### Hierarchical loss of protein-coding genes reflects microRNA losses

To test whether the extensive losses in syndermatan microRNA complements were reflected in other features of their genomes, we compared genome size, genome assembly quality (N50), number of annotated protein-coding genes and protein-coding core gene complements (BUSCO) (Manni et al. 2021). This revealed that genome size estimates or assembly quality could not explain microRNA losses (Figure 2A, R^2^=0.17 and R^2^=0.08, respectively). Instead, we found a correlation between the number of protein-coding genes (numbers taken from (Hagemann et al. 2023)) and microRNA families averaging to 424 protein-coding gene losses per microRNA family loss (R^2^=0.64) (Figure 2B). This suggests that the reduction of protein-coding genes is closely linked to a reduction of microRNA families. To test this further with genes of clear orthology and paralogy, we focused on a highly conserved and universal subset of protein-coding genes (BUSCO: Metazoa node; (Manni et al. 2021). This approach revealed a tight correlation of BUSCO gene and microRNA losses (R^2^=0.93; Fig. 2D), with 11 BUSCO gene losses per microRNA family loss. Mapping the losses on the branches of the most recent syndermatan phylogenetic tree (Vasilikopoulos et al. 2024) suggests a hierarchical and progressive reduction of universal single-copy metazoan orthologs from free-living to epibiotic and endoparasitic ancestors (Figure 2C, D). Thus, we observed 54 shared losses in all syndermatans and additional 61 and 165 shared losses in Hemirotifera (Bdelloidea plus Pararotatoria) and Pararotatoria (Seisonidea plus Acanthocephala), respectively. *Seison nebaliae* lacked 76 and the Acanthocephala 148 further BUSCO genes. The extent of losses in the epibiotic seisonid and both acanthocephalans were reminiscent of previous reports on BUSCO losses in the endoparasitic nematomorphs (Cunha et al. 2023). However, with ∼40% missing metazoan BUSCO genes in *Seison* and Acanthocephala, the extent of losses is highest for all animals studied so far.

**Figure 2:**
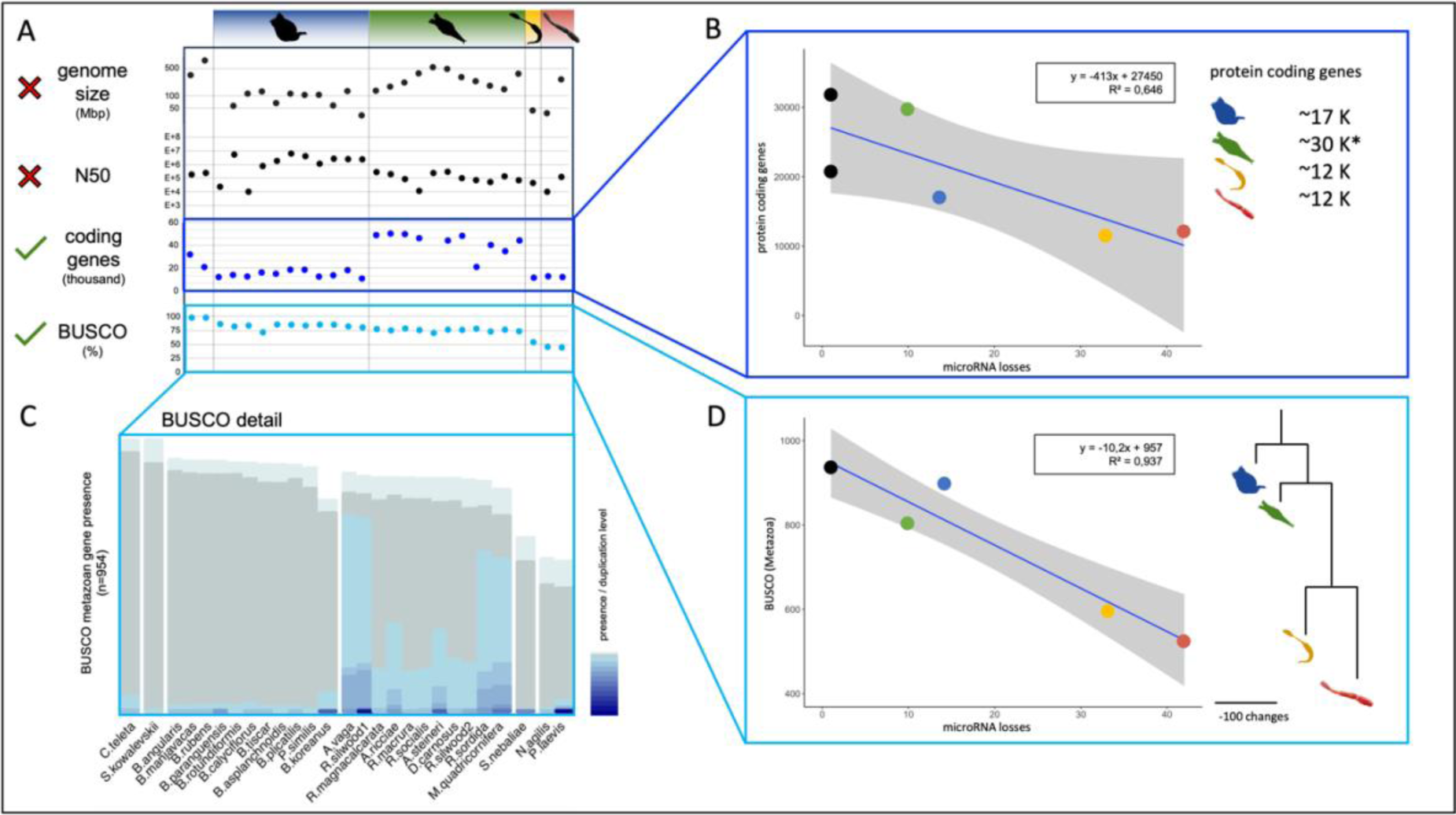
Number of protein-coding genes and protein-coding core gene complements (BUSCO), but not genome size or assembly quality (N50) correlate with microRNA loss. A) Genome size, N50, number of protein-coding genes and metazoan BUSCO completeness of 27 syndermatan species and two outgroup representatives. B) Correlation of protein-coding gene annotations with microRNA loss in representative syndermatans and two outgroup representatives. Grey area corresponds to 95% confidence interval. The asterisk (*) refers to a high number of paralogues in bdelloids due to genome duplication. C) The BUSCO completeness plot highlights the genome duplication in bdelloids. D) Correlation of metazoan BUSCO core genes with the loss of microRNA genes in representative syndermatans and reconstructed tree of BUSCO gene losses. Grey area in graph corresponds to 95% confidence interval. Branch lengths correspond to the number of losses.

### Strong enrichment of gene regulation ontologies in the lost BUSCO genes

Subsequently, we addressed the functional implications of extreme gene loss in the host-bound representatives of Pararotatoria. For this purpose, we conducted enrichment analyses of Gene Ontology terms in the 280 BUSCO genes missing in Seisonidea and Acanthocephala (Figure 3A). The missing genes were significantly enriched for particular terms of all three major categories (i.e., *Biological Process*, *Cellular Component* and *Molecular Function*) (Figure 3B). For *Cellular Component* and *Molecular Function*, we observed 1 (intracellular protein-containing complex) and 3 (protein binding, identical protein binding, transcription regulator activity) enriched annotations, respectively. Within the *Biological Process* category, we identified 81 enriched GO terms. Of those, the majority was associated with regulatory terms such as *regulation of biological process*, *biological regulation*, *regulation of cellular process* etc. (Figures 3B, 3C & 3D pink arrows). Consultation of the GeneCards database uncovered that more than 50% of the genes missing in Pararotatoria (Seisonidea + Acanthocephala) code for positive effectors of transcription (such as transcription factor 25, Mediator of RNA Pol II subunits – MED19/10/8/7/20), transcription repressors (such as Negative elongation factor – NELFB/C/D), and other DNA-binding factors. The encoded proteins were components of RNA binding proteins and mediator complexes implicated in the basal RNA Pol II transcription machinery. We also observed involvements in cell division and proliferation as exemplified by Cyclin-Q and Cyclin-H. Genes implicated in the regulation of *Initiation of protein synthesis* like Eukaryotic translation initiation factors 2 (eIF2 subunits D, B1, B2, B3, B5) and 3 (eIF3 subunits K, L,G,M) were additionally absent in pararotatorians.

**Figure 3:**
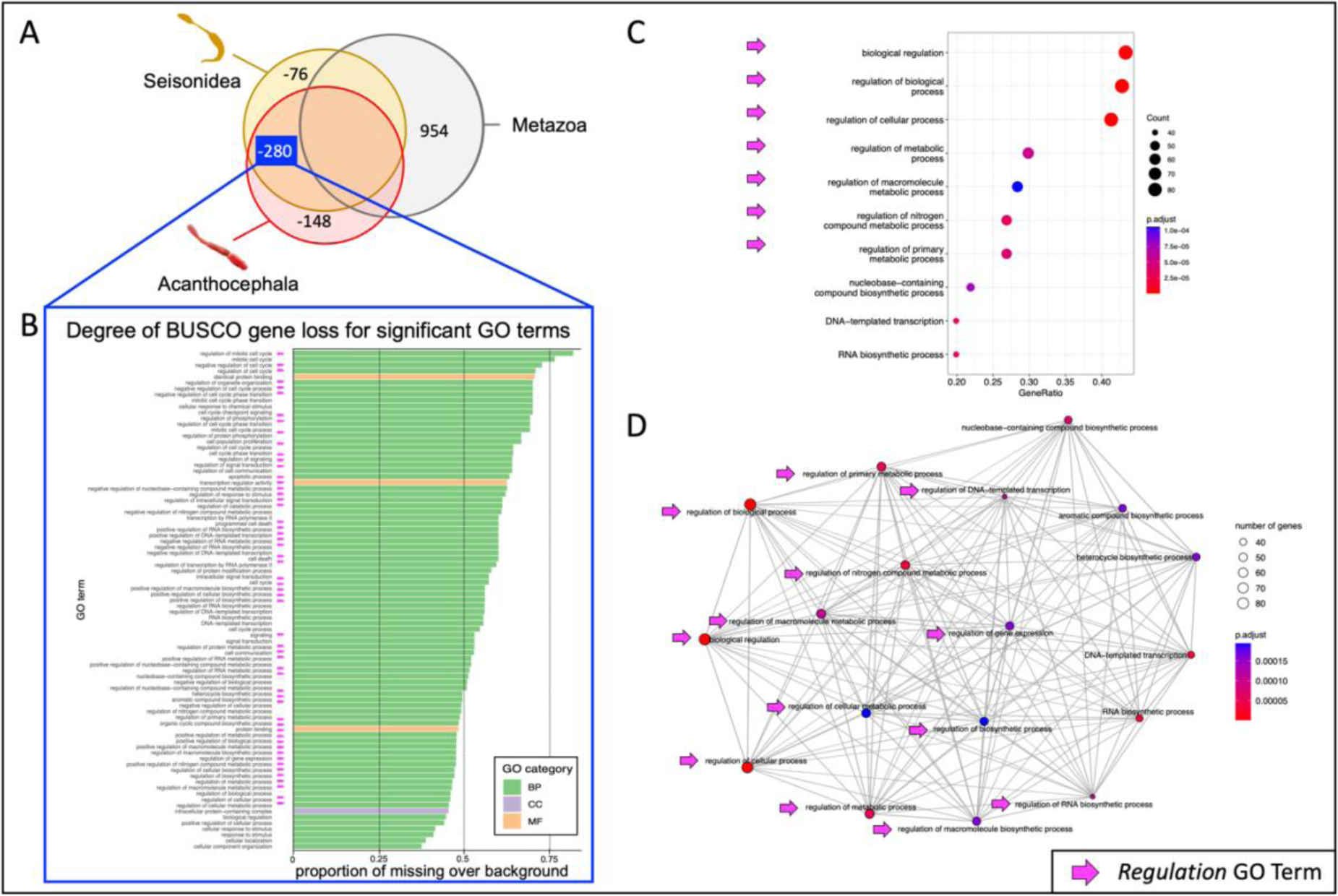
Missing BUSCO genes in Pararotatoria are highly enriched for Biological Regulation functions. A) Venn diagram representing the missing metazoan BUSCO genes in Pararotaria representatives *S. nebaliae* (Seisonidea) and *Neoechinorhynchus agilis* (Acanthocephala). Of the 954 protein-coding genes usually conserved in Metazoa, the seisonid and acanthocephalan collectively lack 280 genes. Additional 76 and 148 genes are absent in Seisonidea and Acanthocephala, respectively. B) Significantly enriched gene ontology (GO) terms in the metazoan BUSCO genes missing in Pararotatoria. Enrichment of 3 *Molecular Function* GOs and one *Cellular Component* GO contrasted with 81 enriched *Biological Process* GO terms. Note the high number of terms referring to regulation (pink arrows). C) Significantly enriched *Biological Process* GOs for missing genes in pararotatorians, whereby each circle represents one term. Color and weight of the dots correspond to the scale on the right. Note the high number of regulatory terms (pink arrows). D) Network analysis of the *Biological Process* GOs. Weight and color of dots is based upon the scale to the right. Note the high number of terms referring to regulatory implications (pink arrows).

### Morphological losses retrace gene and gene regulator losses

To quantitatively assess morphological character losses and gains, we expanded the list of syndermatan entries in the matrix by Deline et al. (Deline et al. 2018), which was derived from Ax ((Ax 2012; Ax 2013a; Ax 2013b) see Materials and Methods). We regarded characters or states present in the gnathiferan or syndermatan last common ancestors as plesiomorphies. In contrast, we defined characters or states that should have arisen on individual branches of the syndermatan tree as apomorphies. The resulting character matrix (Figure 4A) led to trees giving plesiomorphy losses and apomorphy gains (Figure 4B). These illustrated the most extensive loss of plesiomorphic characters such as trunk and head ciliation, the total digestive system, and protonephridia in acanthocephalan evolution (see Figure 4C for details). Notably, branch length relations were highly similar in trees depicting losses of plesiomorphic characters (Figure 4B), microRNA families (Figure 1B) and BUSCO genes (Figure 2B). In line with this, we observed strong associations of plesiomorphy and microRNA losses (Figure 4D) on the one hand, and BUSCO genes and plesiomorphic character losses on the other (Figure 4E) (R^2^=0.72 in both cases).

**Figure 4:**
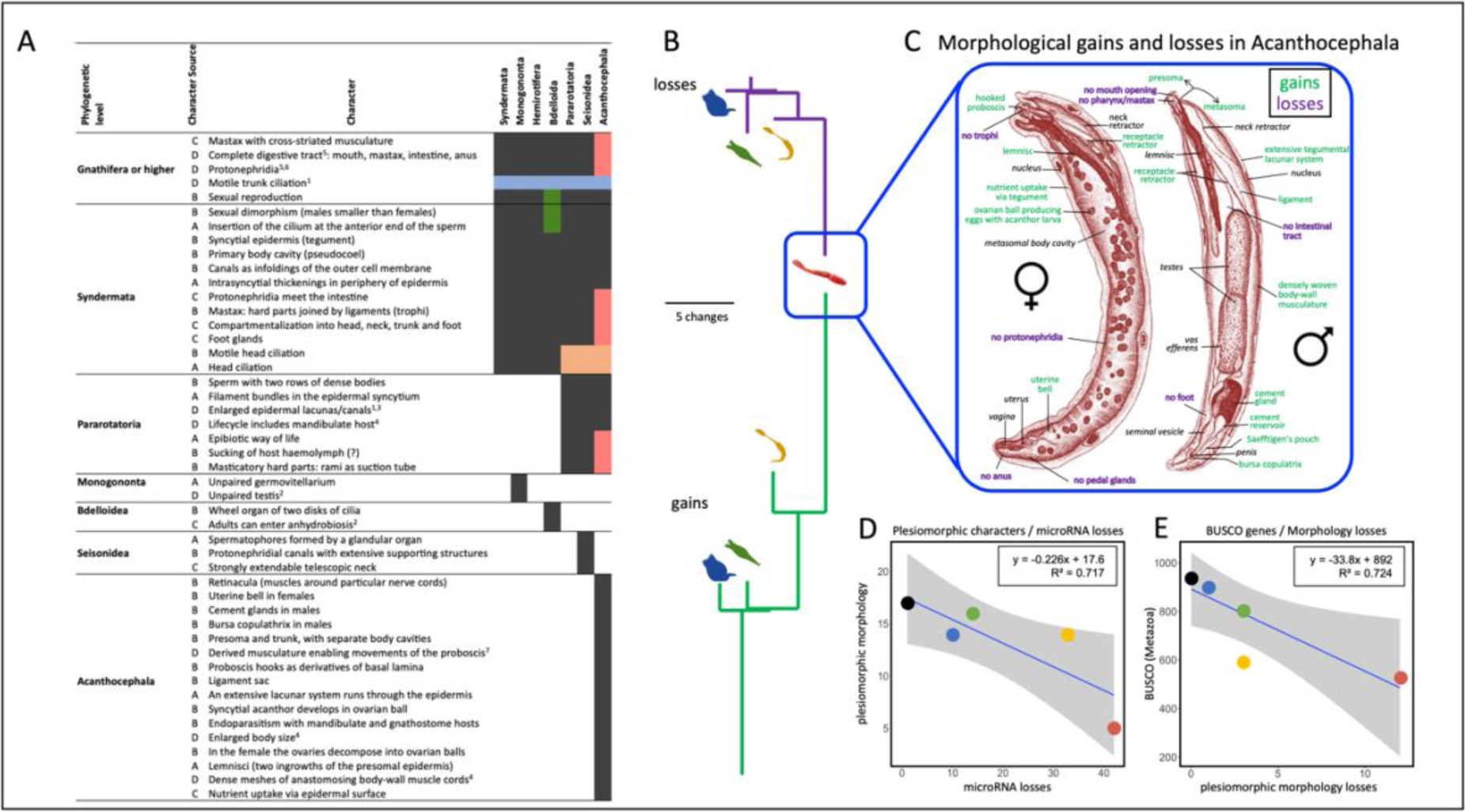
A) Syndermatan character matrix combining novel characters with the ones compiled by Deline et al. (Deline et al. 2018). Dark gray vs. white in taxon columns indicates character presence vs. absence. Blue-, red-, green- and orange-colored fields represent absences interpreted as losses. Column 2 indicates the source of respective characters: A - Deline et al. (Deline et al. 2018), B - modified wording of character definition by Deline et al. (Deline et al. 2018), C - Ahlrichs (Ahlrichs 1995), D - assessment by authors considering: ^1^Ahlrichs ((Ahlrichs 1997): Fig. 6), ^2^Fontaneto & De Smet (Fontaneto and De Smet 2015), ^3^Nicholas & Mercer (Nicholas and Mercer 1965), ^4^Herlyn (Herlyn 2021), ^5^Ahlrichs (Ahlrichs 1995), ^6^Amin (Amin 1987), ^7^Herlyn & Taraschewski (Herlyn and Taraschewski 2017). B) Schematic trees of the four syndermatan groups representing loss of plesiomorphic (purple, top) and gain of apomorphic (green, bottom) morphological features. Branch lengths correspond to gains and losses. C) Morphological character loss (purple) and gain (green) in the extreme example of Acanthocephala. The eponymous attachment organ (proboscis) is inverted in the juvenile female and everted in the adult male. Drawings refer to *Neoechinorhynchus rutili*, a close relative of *N. agilis*. Modified from Steinsträsser (Steinsträsser 1936). D) Dependence of plesiomorphic morphological characters on loss of microRNA genes in representative syndermatans and two outgroup representatives (same species as in Figure 2). E) Dependence of metazoan BUSCO core genes on the loss of plesiomorphic morphological characters in representative syndermatans and two outgroup representatives. Outgroup species are the same as in Figure 2 and were set to no morphological losses. The last common ancestor of extant Acanthocephala might still have possessed protonephridia. In any case, the two acanthocephalans studied (*P. laevis* and *N. agilis*) lack protonephridia.

In summary, the loss of ∼400 protein-coding genes (Figure 2B, R^2^=0.64) or ∼10 metazoan core genes (Figure 2D, R^2^=0.93) went along with one microRNA family loss; and the loss of ∼4 microRNA families or ∼34 metazoan core genes associated with one lost plesiomorphic morphological feature. Nevertheless, morphological character gains also occurred, and they exceeded amounts of losses on most branches. We noticed the strongest extend of character gains (absolute and relative to losses) for the acanthocephalan branch, followed by the branches to epibiotic seisonids and free-living rotifers. The detailed nature of these new morphological feature (composition of cell types, or gene regulatory networks) remains to be studied, but is unlikely to be driven by the expansion of microRNAs and, hence, new cell types.

### Syndermata retain piRNAs

Previously, loss of Piwi-interacting small RNAs (piRNAs) and DNA methylation was observed in parasitic flatworms and many parasitic nematodes (Tsai et al. 2013; Zheng 2013; Skinner et al. 2014; Sarkies et al. 2015; Fontenla et al. 2021; Sarkies 2022). The loss of piRNAs, in particular, was suggested to be connected to the loss of complexity in parasites (Tsai et al. 2013; Fontenla et al. 2021). However, we found piRNAs and PIWI orthologues in all syndermatans, including the epibiotic and parasitic pararotatorians (Figure 5A-D). In all syndermatan genomes studied, piRNA coding sequences were more densely packed than expected by chance, indicating possible piRNA clusters such as in *Drosophila* and mammals. The *P. laevis* genome assembly exhibited the least compact clustering of piRNA loci (Figure 5C, E). Moreover, we noticed that piRNAs lacked the ping-pong signature in the bdelloid *A. vaga*, whilst being retained in the other syndermatan representatives. Furthermore, there was no indication of genes encoding DNA methyltransferase 1 and 3 (*Dnmt1* and *Dnmt3*) in any of the syndermatans analyzed (Figure 5D). Furthermore, the gene coding for RNA-dependent RNA polymerase (*Rdrp*) was absent in pararotatorians whereas we identified multiple copies in the monogonont (*B. koreanus*) and bdelloid included (*A. vaga*). Curiously, there was evidence for zucchini gene (*Zuc*) presence in *P. laevis* while no counterparts seemed to exist in the other syndermatans, including the acanthocephalan *N. agilis*.

**Figure 5:**
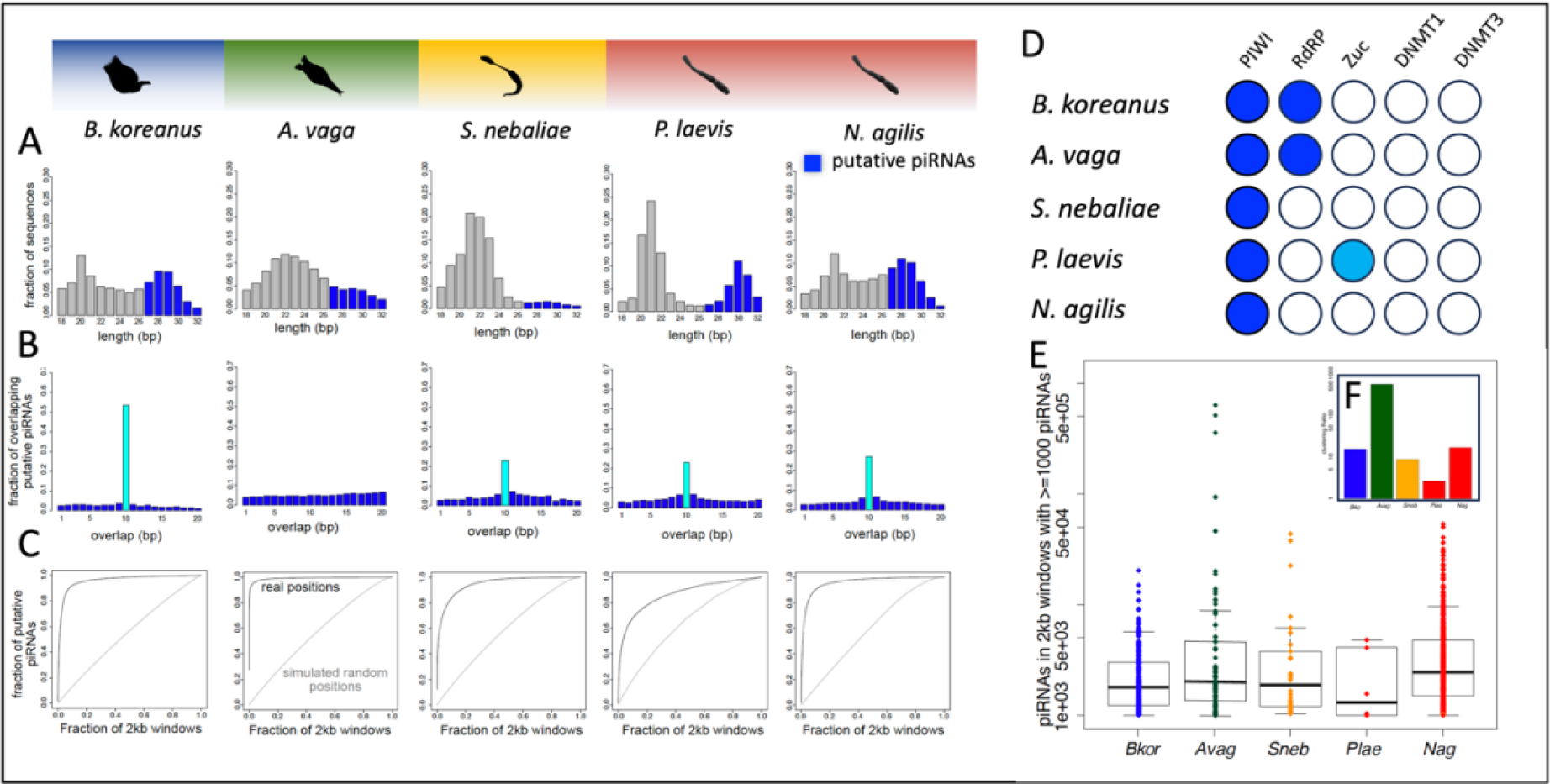
piRNA retention in syndermatans. A) Length distribution of genome-mapped small non-coding RNAs, with putative piRNAs highlighted. B) Distribution of overlap between putative piRNAs on opposite strands, highlighting a peak at 10nt where present, indicative of ping-pong biogenesis. C) Cumulative frequency plot showing the tendency of putative piRNAs to be clustered together along the genome, indicative of potential piRNA-generating loci or piRNA clusters. D) Conservation of proteins related to small non-coding RNA pathways. Annotation of the presence or absence of homologue genes on the basis of best reciprocal blast searches is illustrated for each species accordingly (dark blue - multiple hits, light blue – one hit). E) The piRNAs/2kb windows for each of the 2kb windows with at least 1000 piRNAs mapped. F) The clustering ratio, defined as the ratio of the fraction of the genome containing 90% of randomly shuffled piRNAs compared to the fraction containing 90% of true piRNA positions. A higher clustering ratio thus indicates that a smaller fraction of the genome contains 90% of the piRNAs than expected by chance. *Avag*, *A. vaga*; *Bkor*, *Brachionus koreanus*; *Nag*, *N. agilis*; *Plae*, *P. laevis*; *Sneb*, *S. nebaliae*.

## Discussion

A number of phylogenetic and genomics studies revolutionized our understanding of parasite evolution by largely resolving their relationships to free-living organisms (Aboobaker and Blaxter 2003; Helm et al. 2012; Tsai et al. 2013; Hahn et al. 2014; Foox and Siddall 2015; Sarkies et al. 2015; Lu et al. 2017; Lu et al. 2019; Brabec et al. 2023). Collectively, these investigations demonstrated that morphological reduction is common in the evolution of parasitic life forms. Such simplification may be accompanied by the loss of regulatory microRNAs, which have been proposed to be central drivers of organismic complexity (Sempere et al. 2006; Tanzer and Stadler 2006; Grimson et al. 2008; Heimberg et al. 2008; Peterson et al. 2009; Wheeler et al. 2009; Christodoulou et al. 2010; Berezikov 2011; Fromm et al. 2015; Alberti et al. 2018; Deline et al. 2018; Zolotarov et al. 2022).

We tested this prediction in helminths of the taxon Syndermata, in which lifestyles range from free-living to endoparasitic (Fontaneto and de Smet 2015; Herlyn 2021). Combining novel short non-coding sequencing data with protein-coding transcriptome, genomic and morphological data, we have shown that reduction of morphological features in parasite evolution goes along with loss of microRNAs and protein-coding genes. We observed ∼4 microRNA losses per ∼34 lost metazoan core genes and one lost morphological feature in Syndermata. Of note, we detected losses of traits, core genes and microRNAs on all branches of the syndermatan tree, with the extent accumulating in the lineage to acanthocephalans. This supports that genomic regression of parasites is a strong underlying and hierarchical process (see (Jackson 2015; Adams et al. 2020)) and not just sporadic or mosaic (Tsai et al. 2013; Hahn et al. 2014; Zarowiecki and Berriman 2015; International Helminth Genomes Consortium 2019). Our results additionally complement previous evidence for a significant role of microRNA losses in parasite evolution in neodermatan platyhelminths (Fromm et al. 2013). Thus, both, parasitic syndermatans (pararotatorians) and neodermatans, have lost the majority of their microRNA genes present in their free-living ancestors (contra (Jackson 2015)). However, the extent of losses of widely conserved microRNAs as observed here in seisonid (67%) and acanthocephalan (up to 85%) syndermatans clearly exceeds the corresponding maximum observed in parasitic flatworms (50%) (Fromm et al. 2013). With only 12 microRNA genes in total, the acanthocephalan *P. laevis* has the lowest number of microRNAs reported for any macroscopic bilaterian animal to date. Also, gains of novel microRNAs occurred in all syndermatan species, but their amount was comparatively low for the parasites.

Six novel microRNA families might further substantiate syndermatan monophyly, while numbers of shared losses (9 losses for Syndermata, 2 additional losses in Hemirotifera, and 16 more losses in Pararotatoria) accord with the most recent phylogenetic analysis (e.g. (Vasilikopoulos et al. 2024)). In any case, the hierarchical pattern of microRNAs loss clearly supports a ‘degressive’ evolution scenario, but also follows Dollo’s Law of the irreversibility of evolution (Dollo 1893). Despite their extreme microRNA losses, it is therefore still possible to taxonomically diagnose acanthocephalans based on their minimal retention of microRNAs as syndermatans as they possess the one eumetazoan microRNA family (MIR-10), one bilaterian microRNA (LET-7), two protostome microRNA families (BANTAM and MIR-750) as well as one microRNA family shared only with the other rotifers (Figure 1) (see also (Fromm 2024)).

Given their functional implication in gene regulation (Bartel 2018), the loss of microRNAs might reflect either reduced regulation of preserved protein-coding genes or loss of these targets. Limited protein-coding gene losses are common in free-living invertebrate (Steinworth et al. 2023) and vertebrate groups (Emerling et al. 2017) and are considered important in the evolution of metazoans (Albalat and Cañestro 2016; Fernández and Gabaldón 2020; Guijarro-Clarke et al. 2020). In the present study, confident resolution of orthology-paralogy relations amongst the assumedly 18 000 - 20 000 metazoan protein-coding genes (Erwin 2009) was impaired by overall high divergence levels and genome annotation qualities (Figure 2B). Thus, we focused on highly conserved and single-copy metazoan core genes (BUSCO (Manni et al. 2021)) as proxies for the study of gene repertoire evolution in Syndermata. Our analyses of 954 metazoan core genes revealed significant losses in syndermatans that clearly mirrored microRNA losses, and with ∼40%, losses in Seisonidea and Acanthocephala were higher than reported for any bilaterian before (Figure 2). GO enrichment analysis of the core genes missing in host-bound seisonid and acanthocephalan species revealed a strong enrichment of regulatory terms. This further supports a gene regulatory decline in the pararotatorian stem line through microRNA loss (Figure 3).

Altogether, a scenario seems likely according to which microRNAs are lost upon the loss of their targets, the protein-coding genes. These losses, in turn, seem to have affected ancestral morphological features of the species as evidenced by strong negative correlations of gene losses with reductions of syndermatan basal pattern traits (Figure 4).

However, while we observed very limited gain of novel microRNAs in each species and are unable to reliably identify novel protein-coding genes, there was substantial gain of morphological novelties in acanthocephalan evolution. As for now, the genetic and cellular nature of these new morphological feature is not clear and more detailed analyses in the other groups might reveal novel features in them, too Nevertheless, present evidence corroborates that parasite evolution is more than just simplification but also includes the differentiation of novel characters. In acanthocephalan evolution, a prominent example for simplification is the loss of the digestive tract while the eponymous hooked attachment organ (proboscis) at the anterior body pole exemplifies a morphological gain (Meyer 1933; Taraschewski 2014; Herlyn and Taraschewski 2017). With these and other evolutionary changes, acanthocephalans have very successfully mastered the challenges of a parasitic lifestyle. In fact, new species are frequently being described upon in-depth investigation of gnathostome vertebrates which at least occasionally feed on infected mandibulate arthropods (Amin 2024; Olivera and Campião 2024). Thus, the previous inventory of about 1200 described species of Acanthocephala (Gibson et al. 2014) is probably far below the actual diversity.

Notably, acanthocephalans and the other syndermatans studied possess the piRNA machinery and express piRNAs. This finding shows that there is no general absence of the piRNA pathway in parasites (Tsai et al. 2013; Fontenla et al. 2021), and other factors beyond parasitism may be the actual drivers of piRNAs loss from certain lineages (Sarkies 2024). It is further worthwhile mentioning that piRNAs retained the characteristic ping-pong signature in most syndermatans studied, except for the bdelloid *A. vaga*. Potentially, the ping-pong pathway that amplifies piRNAs in other species (Brennecke et al. 2007) may not be required in *A. vaga*. Here, loss of ping-pong might be compensated by multiple *Rdrp* copies ((Nowell et al. 2021); present study) as suggested by ping-pong replacement by RDRP in the nematode *C. elegans* (Batista et al. 2008). Still, absent genes for Dnmt1 and Dnmt3 suggest reduced DNA methylation ability in syndermatans, which herein correspond with other helminth taxa.

In conclusion, we herein report evidence for an association between the loss of morphological complexity, the loss of protein coding genes and the loss of microRNAs in syndermatans. While we also observe *novel* traits, these remain to be analyzed for their genetic basis and whether they are true novelties or rearrangements of existing parts (e.g. cell types). Our findings prompt further investigation of the mechanisms whereby loss of microRNAs might cause loss of complexity and whether microRNAs “follow” the loss of their protein-coding targets, or if their loss comes first. For this, a thorough characterization and quantification of cell types in each species as well as high quality annotations of protein-coding orthologues between the species and the occurrence of microRNA target sites within their 3’UTRs would be an exciting next step. The other main regulatory elements of metazoan genomes, Transcription factors or RNA binding protein have herein not been studied, as their Metazoan wide annotation poses a significant curational challenge, and hence their role in reduced complexity or the evolution of complexity in general, remains unknown.

The approximation of animal complexity by morphological characters has drawbacks as it often remains unclear whether new characters represent rearranged previously existing features or true novelties, i.e. by the evolution of new cell types. Single cell RNAseq methods hold great promise for the future, but currently lack standardized methodologies for comparative studies of whole adult animals at scale. Such data will further enable to confirm whether the loss of coding and non-coding genes in parasite evolution is a common feature in all Metazoans.

## Material and Methods

### Sample collection

About 1130 males and females of *S. nebaliae* (Seisonidea) were collected in 2018-2019 in agreement with the Centre National de la Recherche Scientifique from the surface of opossum shrimps (*Nebalia bipes*) gathered at low tide from rock pools in the tidal flats off Roscoff (France, Brittany). For RNA extraction, 33 males and females of *N. agilis* (Acanthocephala: Eoacanthocephala) were excised from the intestines of thin-lipped mullets (*Chelon ramada*) captured in 2021 in Adriatic coastal waters on behalf of the Po Delta Park administration. Altogether, 35 males and females of *P. laevis* (Acanthocephala: Palaeacanthocephala) were collected from host intestines (common barbel: *Barbus barbus*, European eel: *Anguilla anguilla*) caught by authorized fishermen in a gravel pit near Gimbsheim (Germany) and in the Weser River near Gieselwerder (Germany) in 2006 and 2014, respectively. RNAs from specimen pools of *S. nebaliae* and *N. agilis* were extracted with TriReagent (Invitrogen). The *P. laevis* specimens were distributed over three samples from which RNAs were extracted using TriReagent or miRVana (Thermo Fisher Scientific).

### Genomes

Available genomes for Monogononta (Kim et al. 2018; Blommaert et al. 2019; Park et al. 2020; Byeon et al. 2021; Kim et al. 2022), Bdelloidea (Nowell et al. 2018; Simion et al. 2021), *S. nebaliae* (Mauer et al. 2021) and Acanthocephala (Mauer et al. 2020) and the genomes of outgroup species *Capitella teleta* and *Saccoglossus kowalevskii* were downloaded from Genbank. gDNA reads and nuclear draft genome of N. agilis can be retrieved from Genbank under BioProject accession PRJNA1126620 and BioSample accession SAMN41948513. Details on generation and key features of the draft genome are given elsewhere (Supplementary Note S2 of (Schmidt et al. 2022))

### Next-Generation Sequencing of short RNA

For *S. nebaliae* and *N. agilis* one smallRNA sequencing dataset and for *P. laevis* three smallRNA sequencing datasets were generated and sequenced using illumina by a commercial supplier and the Norwegian Sequencing Centre in Oslo. smallRNA sequencing data is available under BioProject PRJNA1127840.

### Small RNA processing and microRNA prediction

All 26 genomes were subject to MirMachine analyses using protostome models and lophotrochozoan nodes (Umu et al. 2023) to arrive at a set of conserved microRNA families. In the cases of *B. koreanus* (SRR19792092, SRR19792093, SRR19792094) and *A. vaga* (SRR3185495 & SRR3187155), publicly available smallRNAseq data was used with MirMiner. For *S. nebaliae*, *N. agilis* and *P. laevis* own smallRNAseq data was produced (see above). All smallRNAseq datasets were processed with miRTrace (Kang et al. 2018) and analyzed in MirMiner (Wheeler et al. 2009) for MirMachine confirmation and the prediction of novel or species-specific microRNAs. Novel microRNAs were blasted to genomes without smallRNAseq data available. All microRNA complements are available in MirGeneDB.org (Fromm et al. 2022).

### piRNA prediction and analyses

Small non-coding RNAs were aligned to the genome using Bowtie (Langmead et al. 2009), to produce sam files. These were processed using samtools (Li et al. 2009) and bedtools (Quinlan and Hall 2010) to generate bed files. The bed files were read into R and processed to determine the length of reads, to test whether there were any sequences with typical length associated with piRNAs (27nt or longer up to a maximum of 32 base pairs). Reads with >=27 base pairs that mapped to opposite strands were identified and the overlap between them tabulated to check for the signature of ping-pong biogenesis, i.e. a 10 nucleotide overlap (Brennecke et al. 2007). To test for the presence of potential piRNA clusters, the number of predicted piRNAs (>=27 base pairs) in each 2kb window across the genome was extracted. The windows were sorted by decreasing piRNA density and the cumulative fraction of the total calculated as increasing numbers of windows were added. Clustering of piRNA loci was interpreted as indicating that some regions have very high density, leading to a curve that rapidly plateaus, compared to the random expectation where the piRNA positions were shuffled across the genome and the relationship between the total fraction of piRNAs and the number of windows is more linear (Beltran et al. 2019). From this, the clustering ratio was defined as the number of windows needed to cover 90% of the piRNA sequences when positions were randomized divided by the number of windows needed to cover 90% of piRNA sequences in their real positions. Higher clustering ratio thus indicates that a larger proportion of the piRNAs are in a smaller number of windows. R code for the analysis is available at Github: https://github.com/SarkiesLab/piRNA_rotifer.

### BUSCO and functional enrichment

All genomes were analyzed with BUSCO v5.4.3 using the Metazoa node (954 genes) (Manni et al. 2021). The IDs were used to obtain Gene Ontology (GO) terms from OrthoDB v10 database (Kriventseva et al. 2019). We then performed an enrichment analysis using the 280 genes missing from both *S. nebaliae* and *N. agilis* as the gene set, and the entire metazoan dataset as the backbone. The over-representation analysis was done for all three GO categories (Biological Process, Cellular Component, Molecular Function) using clusterProfiler package (Yu et al. 2012; Wu et al. 2021) in R (Team 2013). The computations were performed on resources provided by Sigma2 - the National Infrastructure for High-Performance Computing and Data Storage in Norway.

### Morphological character analyses

We created a binary matrix of morphological characters of Syndermata starting from the phylogenetically broader compilation by Deline et al. (Deline et al. 2018), which in turn referred to characters in three text books by Ax (Ax 2012; Ax 2013a; Ax 2013b). In Figure 4 (A) we have indicated when we have adapted character definitions by Deline et al. We have also added characters to our morphological matrix. References to adaptations and additions are given for each character in the legend to that table (Nicholas and Mercer 1965; Amin 1987; Ahlrichs 1995; Fontaneto and De Smet 2015; Herlyn and Taraschewski 2017; Herlyn 2021).

## Supporting information

suppFile

## Acknowledgment

We acknowledge funding through the Tromsø forskningsstiftelse grant (TFS) [20_SG_BF ‘MIRevolution’] to BF and contributory funding by German Research Foundation (Deutsche Forschungsgemeinschaft, DFG: HE 3487/5-1) to HH.

